# Lactate metabolism antagonizes antiviral innate immunity by promoting VDAC1-lactylation-mediated mtDNA release

**DOI:** 10.1101/2025.05.09.653027

**Authors:** Yi Wu, Lu Li, Yiyi Liu, Shenyuan Wang, Fanhua Meng, Tao Li, Hongmei Xiao, Fang Wan, Yingchun Liu, Bing Han, Huanmin Zhou, Chunxia Liu, Yanru Zhang, Junwei Cao

## Abstract

Foot-and-mouth disease (FMD) is an acute, highly contagious disease caused by the foot-and-mouth disease virus (FMDV). Due to its zoonotic nature, this disease not only significantly impacts livestock productivity but also poses a serious threat to human health. FMDV exhibits distinct host specificity, mainly infecting cloven-hoofed animals such as cattle, sheep, pigs, and deer, while perissodactyl animals like horses and donkeys show natural resistance. Elucidating the interspecies differences in response to FMDV infection is essential for understanding the pathogenesis of the virus. In this study, we report that FMDV infection in horse epithelial cells induces the production of lactate, which modulates the function of VDAC1 protein through lactylation, facilitating the release of mtDNA from mitochondria into the cytoplasm. This process activates the cGAS-STING signaling pathway, which stimulates interferon-β expression and suppresses FMDV replication. Our findings provide new insights into the mechanism behind horse resistance to FMDV and lay a theoretical foundation for further research concerning the differential pathogenesis of FMDV infection across species.

## Introduction

Foot-and-mouth disease virus (FMDV), the first animal virus ever discovered, is a small RNA virus belonging to the family *Picornaviridae* and the genus *Aphthovirus*. Its genome consists of a single positive-strand RNA of approximately 8,400 nucleotides, encapsulated in an icosahedral capsid (Jamal and Belsham, 2013; Jamal and Belsham, 2018). FMDV exhibits high genetic variability and is classified into seven distinct serotypes: O, A, C, Asia 1, SAT 1, SAT 2, and SAT 3, with no cross-protection between different serotypes(Brito and Rodriguez et al., 2017). FMD affects a wide range of cloven-hoofed animals, including cattle, pigs, sheep, goats, and about 70 species of wild cloven-hoofed animals (Barman and Patil et al., 2020; Dahiya and Subramaniam et al., 2021). Interestingly, equine species such as horses and donkeys are naturally resistant to FMDV infection, but the underlying mechanisms remain unknown.

It was previously believed that the natural resistance of horses to FMDV infection was due to the lack of integrin receptors on their cell membranes that the virus binds to. However, this theory has been challenged by subsequent research. A previous study by our laboratory found that both bovine and equine nasopharyngeal tissues express the integrin receptors, including αvβ1, αvβ3, αvβ6, and αvβ8, all of which are currently recognized as FMDV receptors(Wanfu, 2018). This discovery indicates that horses have the potential to be infected by FMDV through the same receptors as cattle. Other studies have also cloned integrin subunits αv, β3, and β6 from both cattle and horses and introduced them, either individually or in combination, into non-permissive SW480 and COS1 cells. These studies found that FMDV could replicate in SW480 cells transfected with horse β3/β6 integrin subunits and in COS-1 cells transfected with horse αvβ3/αvβ6 integrins, but not in equine kidney epithelial cells (EKECs) (Wang and Mao et al., 2016). These findings suggest that the resistance of horses to FMDV infection is not related to integrin receptor expression, prompting us to re-examine the alternative mechanisms responsible for species-specific resistance.

Protein lactylation is a newly discovered post-translational modification first proposed by Zhang et al. in 2019. This modification is regulated by glycolysis and L-lactate levels (Zhang and Tang et al., 2019). Pyruvate, the precursor of lactate, is transported into cells through monocarboxylate transporters (MCTs). In the cytoplasm, lactate can be oxidized back to pyruvate via the pyruvate dehydrogenase complex (PDH), which irreversibly removes lactate from the system. Alternatively, lactate can be converted to glucose through gluconeogenesis and participate in histone and non-histone protein lactylation. Lactate also acts as a signaling molecule via G protein-coupled receptor 81 (GPR81).

Current research has shown that lactylation is involved in various biological processes, such as glycolysis, macrophage polarization, nervous system regulation, and even grain development in rice. However, whether protein lactylation affects FMDV replication has not yet been explored.

In this study, we established an antibody library against FMDV and applied immunoprecipitation combined with mass spectrometry to explore the shared and distinct mechanisms triggered by FMDV infection in primary cells isolated from both susceptible (cows and goats) and resistant (horses) species. Our study discovered that the FMDV replication is inhibited by protein lactylation. In vitro infection of horse epithelial cells with FMDV induced the lactylation of host protein VDAC1, which subsequently promoted the release of mitochondrial DNA (mtDNA) into the cytoplasm, activating the interferon signaling pathway, and thereby reducing FMDV replication. Our findings revealed that perissodactyls (single-toed or odd-toed animals such as horses and zebras) have evolved distinct antiviral mechanisms, in contrast to artiodactyls (even-toed animals like cattle and goats), by inhibiting FMDV replication through VDAC1 lactylation.

## Results

### Differences in FMDV copy number over time in epithelial cells from cattle, goats, and horses after infection

To investigate whether FMDV can infect horse epithelial cells and replicate therein, we first observed the morphological changes in epithelial cells from the three species following FMDV infection and measured viral copy numbers at different times. The results showed that both bovine and caprine epithelial cells gradually died after FMDV infection, with approximately 80% cell death occurring by 24 hours post-infection (hpi). In contrast, although being infected under the same conditions, no significant morphological changes were observed in horse epithelial cells (Figure 1A), indicating that horse cells may possess resistance or tolerance to FMDV, thus limiting viral replication and avoiding widespread cellular damage.

**Figure 1.**
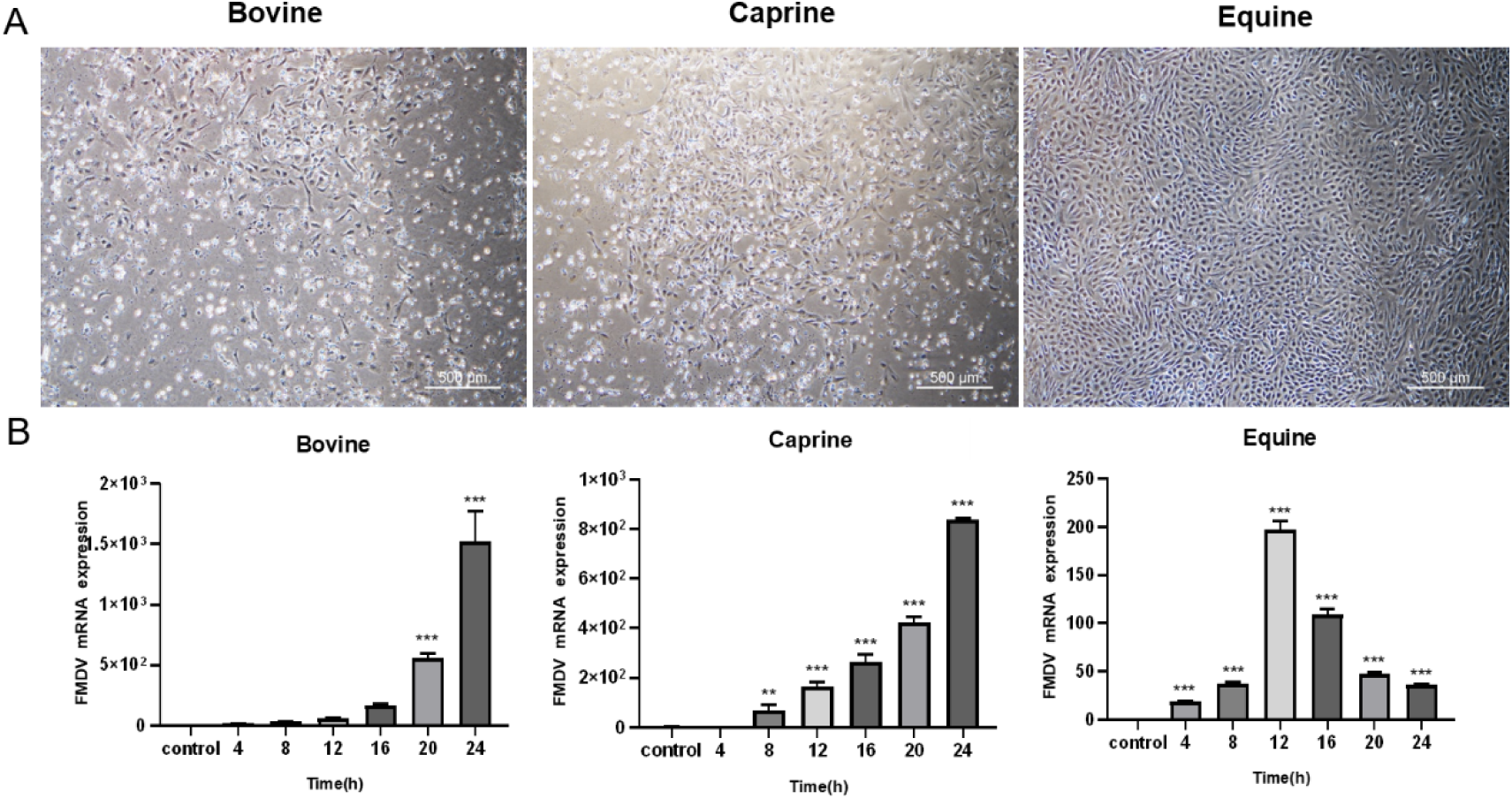
Variation of FMDV copy number in epithelial cells from different species. (A) Morphological changes in nasopharyngeal epithelial cells 24 hours after FMDV infection (400X). (B) Time course of FMDV infection in epithelial cells from different species.

To assess viral replication, we quantified FMDV RNA copy numbers using RT-qPCR (Figure 1B). The results showed that the viral copy numbers increased over time in bovine and caprine epithelial cells. In contrast, in equine epithelial cells, the viral copy numbers remained consistently lower and began to decrease after 12 hpi. These findings suggest that horse cells can also be infected by FMDV but possess certain mechanisms to prevent viral replication, leading to viral clearance beginning around 12 hpi.

### Establishment of an FMDV monoclonal antibody library of hybridomas

To compare the differences in mechanism of FMDV infection between cattle and horses, we created an antibody library against FMDV using proteins from FMDV-infected cells as antigens and employed an antibody microarray. A laser scanner was used to excite and record the fluorescence signals from the chips. The results showed that the internal positive control values fell within the expected range (Figure 2A). The signal-to-noise ratio (S/N), calculated as the ratio of antibody fluorescence intensity to the background signal, was determined to be S/N > 10 (Figure 2B), indicating satisfactory chip quality for subsequent analysis.

**Figure 2.**
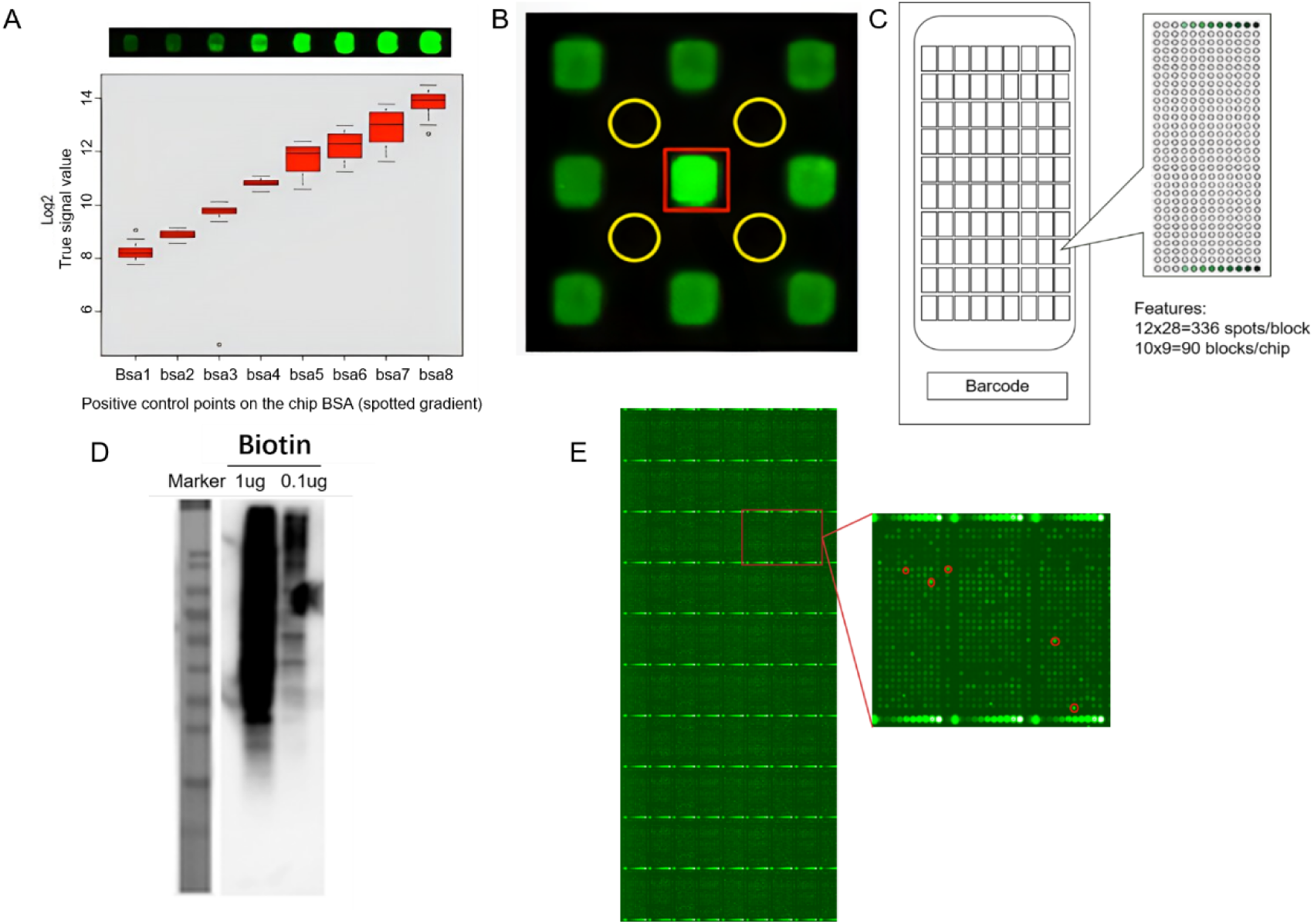
FMDV antibody library preparation. (A) Internal reference signals on the chip. (B) Signal-to-noise ratio of antibody spots: yellow circles - background noise; red squares - antibody fluorescence signals. (C) Diagram of mAb distribution on the chip: green label - different concentrations of the BSA internal reference antibody, darker colors - higher concentrations. (D) Western blot validation of biotin-labeled proteins. (E) CY3 signal map of sample-antibody microarray hybridization.

The antibody chips were arranged in 9×10 square grid, with each chip block containing 336 antibody spots arranged in a 12×28 square array. Each complete chip contained 30,240 antibodies, including 1,080 internal control spots and 29,160 FMDV-targeting antibody spots (Figure 2C). Proteins extracted from FMDV-infected cells were biotinylated, and the labeling efficiency was confirmed by western blot (WB) using two different loading amounts (1 μg and 0.1 μg). The results showed that the biotin-labeled proteins covered the entire lane regardless of the loading amount, indicating successful biotin labeling (Figure 2D). Biotin-labeled protein samples were hybridized with the qualified antibody chips and scanned using a mAbArray laser chip scanner. Antibody spots with significantly different fluorescence signal intensities (*p*<0.05) were identified. We successfully selected five mAbs (Figure 2E) and assessed their antigen affinity by ELISA. The results showed that, among the five mAbs, CL029884 and CL012856 exhibited the highest affinities (Table 1), and were chosen as the best FMDV-specific antibodies for subsequent immunoprecipitation (IP) experiments.

**Table 1.** ELISA measurement of antibody affinity.

| Number<br>Concentration | CL029884 | CL016492 | CL052377 | CL012856 | CL041050 |
| --- | --- | --- | --- | --- | --- |
| 100 ng | 2.683 | 2.258 | 0.774 | 3.398 | 0.555 |
| 10 ng | 1.536 | 0.187 | 0.279 | 2.813 | 0.228 |
| 1 ng | 1.29 | 0.086 | 0.185 | 1.185 | 0.162 |

### Screening of host proteins interacting with FMDV in bovine, caprine and equine nasal epithelial cells

To elucidate the underlying mechanisms of FMDV infection in different hosts, we chose FMDV-infected cattle as the target species, FMDV-infected goats as the positive control, and horses (resistant to FMDV infection) as the negative control. Epithelial cells from these three species were infected with FMDV, followed by immunoprecipitations (IP) using high-affinity monoclonal antibodies (mAbs) selected from our FMDV antibody library. Mass spectrometry (MS) was employed to identify host proteins interacting with the virus. MS analysis revealed that 520, 237, and 236 host proteins interacted with FMDV in bovine, caprine, and equine nasal epithelial cells, respectively (Figure 3A). To better understand the differences in host-virus interactions between susceptible and resistant species, we performed Venn diagram analyses of these identified protein sets. The Venn diagrams revealed 393, 138, and 120 unique host proteins interacting with FMDV in cattle, goats, and horses, respectively. Between the susceptible species, cattle and goats, 40 shared proteins were identified (Figure 3B). These unique proteins likely reflect their distinct response mechanisms to FMDV infection, contributing to the observed differences in susceptibility. Conversely, the shared proteins may represent common host responses to FMDV infection, particularly in susceptible species such as cattle and goats.

**Figure 3.**
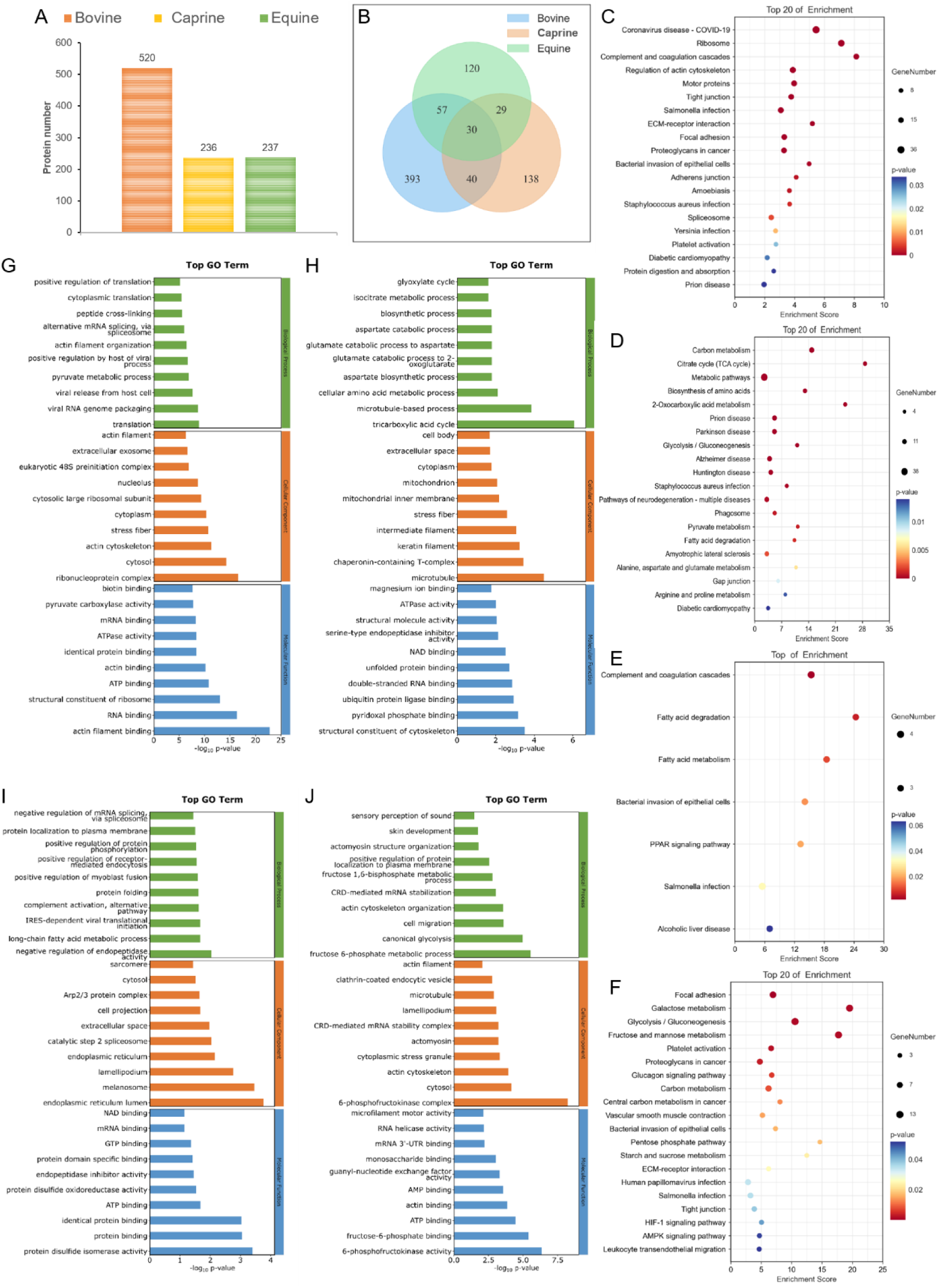
Bioinformatics analysis of host proteins interacting with FMDV. (A) Identification of host proteins interacting with FMDV via mass spectrometry. (B) Venn diagram analysis of host proteins interacting with FMDV. (C-F) KEGG pathway analysis of proteins interacting with FMDV: (C) Bovine-specific proteins; (D) Caprine-specific proteins; (E) Bovine and caprine shared proteins; (F) Equine-specific proteins. (G-J) GO enrichment analysis of proteins interacting with FMDV: (G) Bovine-specific proteins. (H) Caprine-specific proteins. (I) Bovine and caprine shared proteins. (J) Equine-specific proteins.

To further elucidate the mechanisms of these interactions, we performed Gene Ontology (GO) and Kyoto Encyclopedia of Genes and Genomes (KEGG) pathway analyses on the unique and shared proteins sets. KEGG pathway analysis of cattle-specific interacting proteins showed enrichment in pathways such as coronavirus infection, ribosomes, the complement and coagulation cascade, cytoskeletal regulation, and *Salmonella* infection (Figure 3C). GO analysis indicated that these cattle-specific interacting proteins were mainly involved in translation, viral RNA packaging, and virus release from host cells (Figure 3G). Notably, most of the bovine-specific interacting proteins were ribosomes proteins (RPs), such as RPS9, RPL3, RPL10, RPS8, RPL12, RPS6, RPS3A, RPSA, RPL6, RPS14, RPL27A, RPL14, RPS3, RPL13, RPS2, RPL18, RPS27A, RPS11, and RPL17.

KEGG analysis of goat-specific proteins revealed enrichment in pathways associated with metabolism, TCA cycle, prion diseases, Parkinson’s disease, and Huntington’s disease (Figure 3D). GO analysis showed that these proteins were mainly enriched in the tricarboxylic acid cycle, amino acid metabolism, aspartate biosynthesis, and glutamate degradation (Figure 3H).

KEGG analysis of the shared proteins between cattle and goats showed enrichment in complement and coagulation cascades, fatty acid metabolism, the PPAR signaling pathway, *Salmonella* infection, and alcoholic liver disease (Figure 3E). GO analysis showed involvement in negative regulation of endopeptidase, long-chain fatty acid metabolism, IRES-dependent virus translation initiation, and alternative complement activation pathways (Figure 3I). The proteins shared by cattle and goats were more functionally involved in protein processing and the innate immune system. Viruses have evolved strategies to interfere with antiviral immune defenses, including downregulating the complement activation components, C3, C4, and upregulating complement regulatory proteins, as well as recruiting membrane-associated regulatory factors (RCA) that activate complement (Kumar and Kunnakkadan et al., 2020). When bovine and caprine epithelial cells were infected with FMDV, host-virus interactions with C3 and C4 proteins were detected, suggesting that FMDV could also limit the complement response by inactivating C3 and C4, thereby disorganizing the immune system’s attack and promoting viral replication.

KEGG analysis of goat-specific proteins revealed enrichment in pathway associated with prions, Parkinson’s disease, and Huntington’s syndrome, which are all associated with protein misfolding. The health and survival of all organisms depends on correctly folded and functional proteins. Misfolded proteins can become nonfunctional and cause toxic chain reactions when they aggregate. Molecular chaperones play a crucial role in stabilizing and promoting the correct folding of other proteins. Death domain-associated proteins are a class of molecular chaperones that are rich in aspartic acid (D) and glutamic acid (E) and function through their polyD/E regions to promote proper protein folding, prevent protein aggregation, disassemble existing aggregates, and recycle misfolded proteins. GO analysis showed enrichment in the biological processes of aspartate biosynthesis and glutamate metabolism. The sheep-specific interacting proteins also included CCT6A, CCT2, CCT7, CCT4, which are all molecular chaperones. CCTs typically assist cytoskeletal proteins like actin in folding correctly before forming filaments and microtubules(Sternlicht and Farr et al., 1993). Viral infections can alter the normal spatial distribution and biological functions of cellular organelles and macromolecules, with significant changes observed in the cytoskeleton and membrane structures. In our study, GO analysis also showed multiple enriched entries related to the cytoskeleton, indicating that cytoskeletal proteins were affected in goats infected with FMDV. Based on this, we hypothesized that FMDV could affect molecular chaperones in infected goats, causing incorrect folding of cytoskeletal proteins, abnormal structure of filaments and microtubules, and enhanced FMDV replication.

KEGG analysis of equine-specific proteins revealed enrichment in pathways related to adherens junctions, galactose metabolism, glycolysis, gluconeogenesis, fructose and mannose metabolism, platelet activation, and carbon metabolism (Figure 3F). GO analysis showed enrichment in biological processes, such as fructose-6-phosphate metabolic, actin filament organization, glycolysis, and cell migration (Figure 3J). Both KEGG and GO analyses suggested that changes in metabolism were caused by viral infection in horses, and these metabolic changes may be one of the reasons why horses are less susceptible to FMDV.

### Metabolic changes in bovine, caprine and equine nasal epithelial cells before and after FMDV Infection

After discovering that horses infected with FMDV showed functional interactions with specific metabolic proteins, we compared the metabolic changes in epithelial cells from horses, cattle and goats with FMDV infection. First, we determined the changes in glycolysis and oxidative phosphorylation by measuring the extracellular acidification rate (ECAR) and oxygen consumption rate (OCR), respectively. The results showed a significant increase in ECAR following FMDV infection in all three species: cows (*p*< 0.01), goats (*p*< 0.05), and horses (*p*< 0.001) (Figure 4A), indicating enhanced glycolytic activity. In contrast, OCR results showed a significant increase in oxygen consumption in cows and goats, while no significant change was observed in horses (Figure 4B), suggesting that FMDV promoted the oxidative phosphorylation in cows and goats but not in horses. These differences imply that while FMDV infection may promote aerobic respiration in cattle and goats, horses are likely to adopt an alternative metabolic pathway in response to infection.

**Figure 4.**
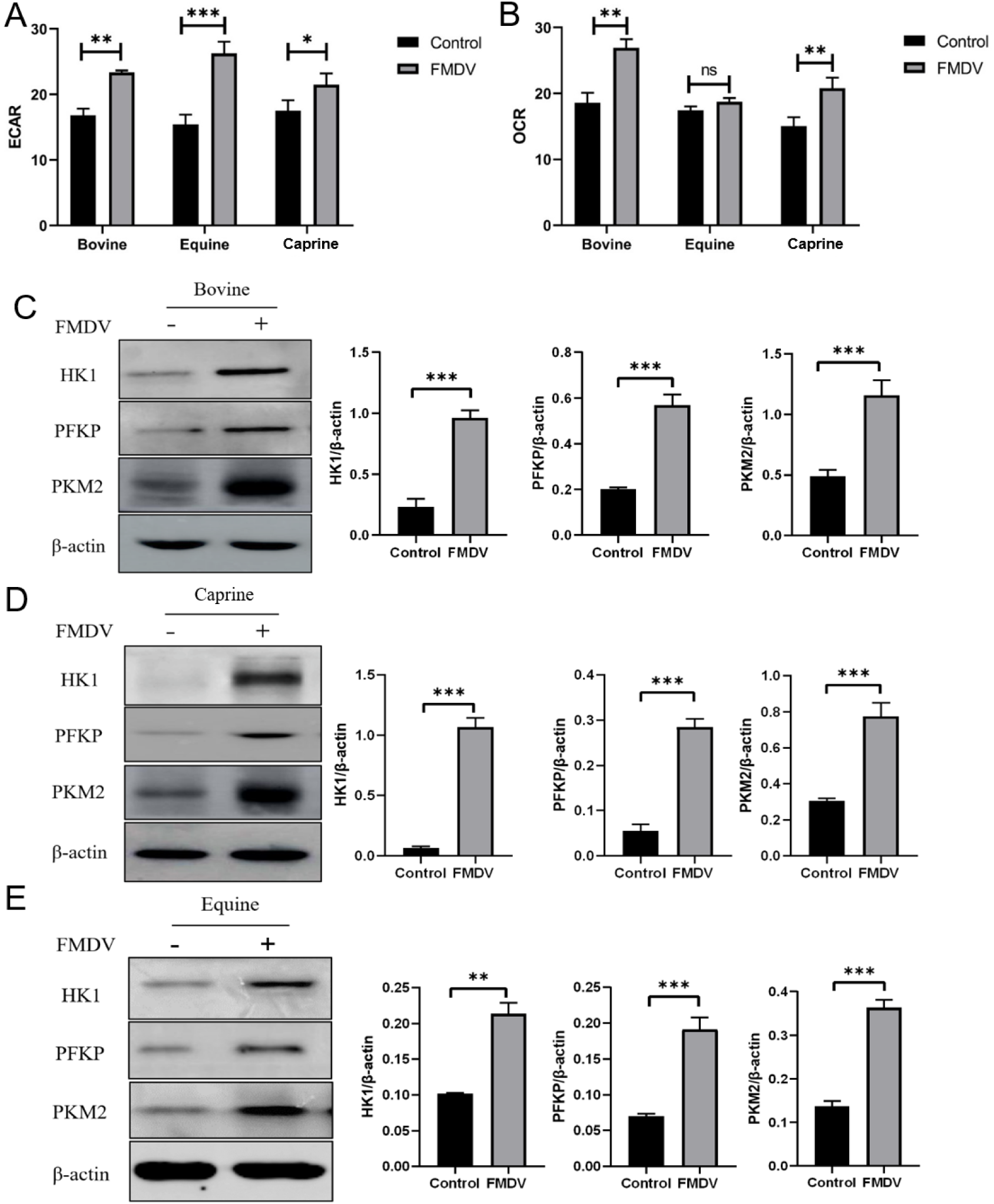
FMDV promotes glycolysis. (A) Extracellular acidification rate (ECAR). (B) Oxygen consumption rate (OCR). (C-E) Western blot assay and quantification of glycolytic enzymes (HK1, PFKP, PKM2) in the three species. Beta–actin was used as internal control for quantification of immunoblot bands. All experiments were repeated at least three times, and values represent means ± SD. Statistical significance was assessed using Student’s *t*-test: \*\**p*< 0.01, \*\*\**p*< 0.001.

To further explore the differences in these metabolic, we measured the expression of three key glycolytic enzymes, HK1, PFKP, and PKM2, which were previously identified as FMDV-interacting proteins specific to horses. Western blot (WB) analyses showed that, after FMDV infection, the expression levels of HK1, PFKP, and PKM2 were significantly upregulated in epithelial cells from all three species at 12 hpi (*p* < 0.001) (Figures C-E), confirming that glycolytic activity was promoted in all three species.

### FMDV infection of nasal epithelial cells from cows, goats and horses results in differences in lactic acid content

Our previous results demonstrated that the expression of three glycolytic proteins (HK1, PFKP, PKM2) was significantly increased by FMDV infection, but no difference observed between horses, cows and goats (Fig. 4C-E). We then focused on pyruvate, the end product of glycolysis, which can be converted into either acetyl-CoA under aerobic conditions or lactic acid under anaerobic conditions. Acetyl-CoA enters the TCA cycle where it participates in oxidative phosphorylation, while lactic acid is generated through the action of lactate dehydrogenase-A (LDHA). Since there was no significant change in the OCR in horses, we hypothesized that pyruvate metabolism might be directed toward increased lactic acid production in horses.

To test this hypothesis, we first measured the LDHA protein expression level in epithelial cells from the three species after FMDV infection. Western blot (WB) analysis revealed a significant increase of LDHA expression in horse compared to the control group (*p*< 0.001) at 12 hpi. In contrast, no significant difference was observed in cattle and goats (Figures 5A-C). Then we directly measured the lactic acid levels and found a significant decrease in cattle (*p*< 0.05) and goats (*p*< 0.001), whereas a significant increase in horses (*p*< 0.05) (Figure 5D). These results indicate that FMDV infection redirects pyruvate metabolism toward lactic acid production in horses but not in cattle or goats.

**Figure 5.**
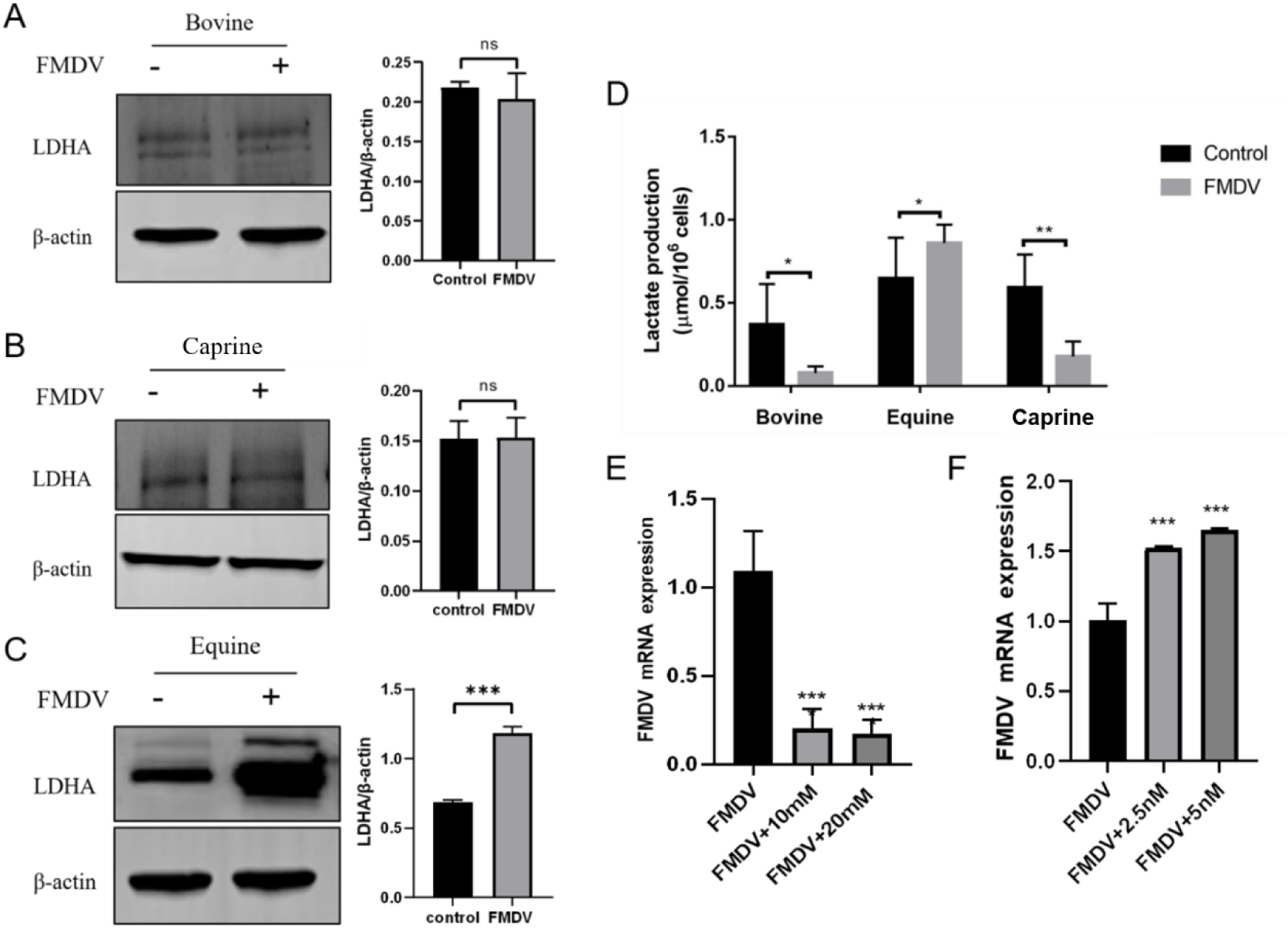
FMDV promotes production of lactic acid in equine epithelial cells. (A-C) WB assay and quantitation of lactate dehydrogenase protein level in cattle, goats and horses after FMDV infection. (D) Lactic acid content after FMDV infection. (E) Effect of exogenous lactic acid addition on FMDV replication. (F) Effect of lactate dehydrogenase inhibitor (GSK2837808A) on FMDV replication. All experiments were repeated at least three times, and values are means ± SD. Statistical significance was assessed using Student’s *t*-test: *ns* > 0.05, \**p* < 0.05, \*\**p* < 0.01, \*\*\**p* < 0.001.

### Effect of lactic acid on FMDV replication

To investigate whether changes in cytosolic lactic acid levels affect FMDV replication, we next conducted experiments using exogenous L-lactate. Previously we tested the effect of lactic acid on cell viability using the CCK-8 viability assay and confirmed that lactic acid ranging from 0 to 40 mM had no significant effect on cell survival. Subsequently, we added exogenous L-lactate at concentrations of 10 mM and 20 mM to FMDV-infected bovine epithelial cells and measured viral copy numbers at 12 hpi. The RT-qPCR results showed a highly significant decrease in FMDV mRNA levels in both 10 mM and 20 mM treatment groups (*p*< 0.001), indicating that lactic acid effectively inhibited FMDV replication (Figure 5E).

We next investigated whether reducing lactic acid production would affect FMDV replication. We treated horse epithelial cells with GSK2837808A, an inhibitor of LDHA, at concentrations of 2.5 nM and 5 nM, and measured FMDV mRNA levels at 12 hpi by RT-qPCR analysis. The results showed a highly significant decrease in FMDV mRNA levels (*p*< 0.001) compared to untreated cells (Figure 5F), suggesting that lactic acid plays a crucial role in suppressing FMDV replication in horse epithelial cells.

### FMDV promotes lactylation of proteins in equine epithelial cells

FMDV was observed to trigger substantial lactate production in equine epithelial cells. Considering the natural resistance of horses to FMDV infection, we were particularly interested in elucidating the potential role of lactate in their antiviral mechanisms and the underlying molecular pathways involved.

In recent years, more and more studies have been conducted on lactate, revealing other functions such as its role as a signaling molecule in biological processes and in epigenetic reprogramming through histone lactylation. Protein lactylation, which was discovered in 2019, has continued to receive much attention, which sparked our interest in determining whether lactylation could affect FMDV replication. Therefore, we performed a proteomic lactylation analysis on FMDV-infected equine epithelial cells with significantly elevated lactate levels. The results revealed a total of 3,889 lactylated peptides, with 4,100 lactylation sites identified on 1,797 proteins (Table 2).

**Table 2.** Comprehensive identification of lactylated proteins in FMDV-Infected equine epithelial cells.

| Title | Number |
| --- | --- |
| 1. Total spectra | 4011701 |
| 2. Peptides | 12308 |
| 3. Modified peptides | 3889 |
| 4. Matched spectra | 449015 |
| 5. Identified sites | 4100 |
| 6. Identified proteins | 1797 |

Notes: **1.** Total spectra: total number of secondary spectra generated by mass spectrometry; **2.** Matched spectra: effective number of spectra matched with theoretical secondary spectra; **3.** Peptides: number of peptide sequences resolved from the matching results; **4.** Modified peptides: number of modified peptide sequences resolved from the matching results; **5.** Identified Sites: number of lactylation sites identified on the identified proteins **6.** Identified proteins: number of proteins resolved from specific peptide sequences.

We performed GO classification and KEGG pathway enrichment analysis on the identified lactylated proteins, calculating *p*-values using Fisher’s exact test to determine if there were significant enrichments in certain functional categories. The GO analysis results showed that in terms of biological processes, the proteins were mainly involved in protein complex oligomerization, oxidative phosphorylation, and the metabolism of purine nucleotides, monocarboxylic acids, ATP, and ribonucleoside triphosphate. In terms of cellular components, the proteins were primarily located in mitochondria, mitochondrial cristae, neuronal projections, the mitochondrial matrix, myelin sheath, and mitochondrial membranes. In terms of molecular function, the proteins were mainly enriched in oxidoreductase activity and in the binding of small molecules, nucleotide phosphates, nucleotides, anions, and protein complexes (Figure 6A). KEGG analysis revealed that the lactylated proteins were enriched in several secondary pathways, including transport and catabolism, cell growth and death, signal transduction, translation, protein folding, protein sorting and degradation, carbon metabolism, endocrine systems, and the immune system (Figure 6B).

**Figure 6.**
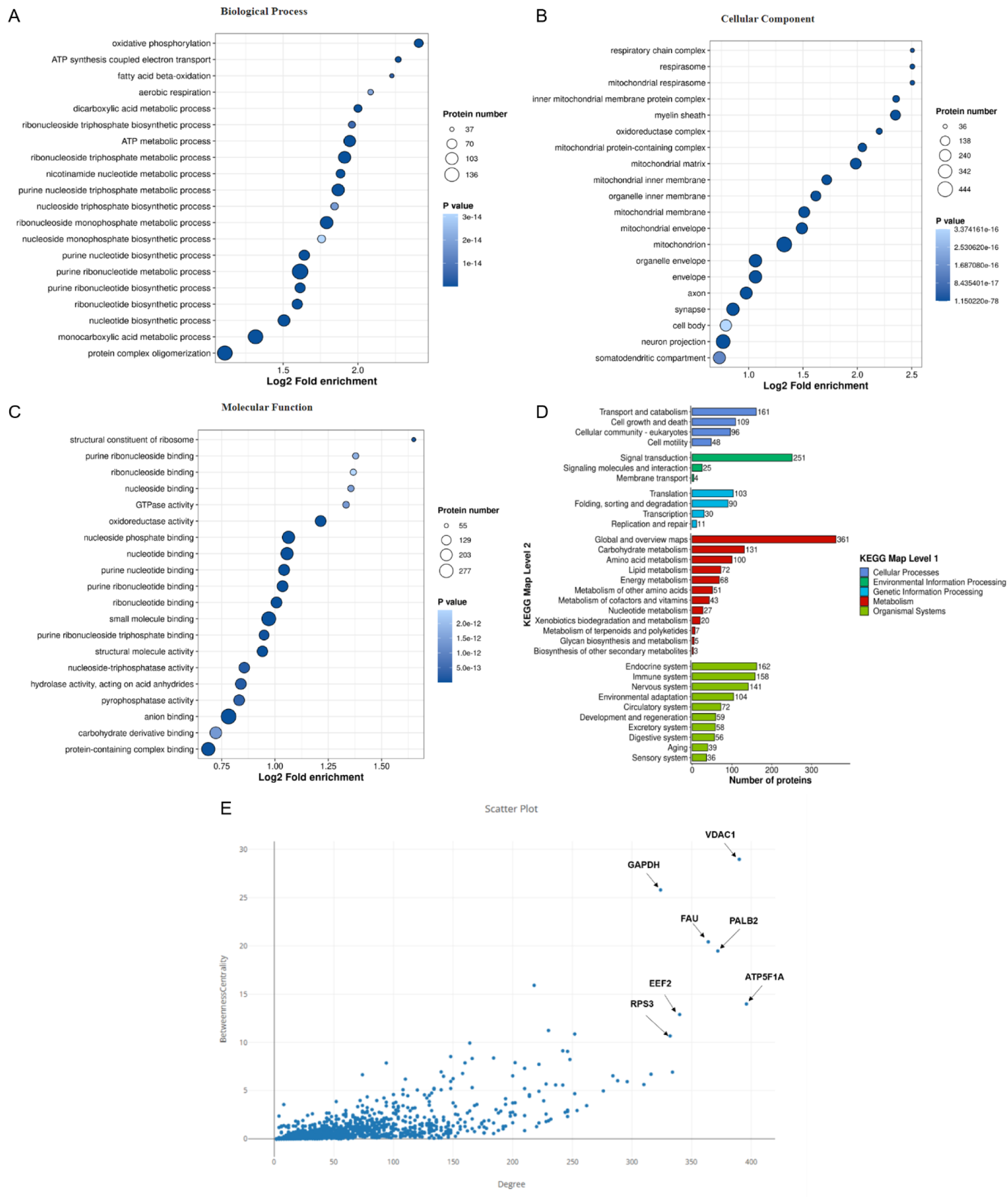
Proteomics analysis of protein lactylation modifications. (A) GO analysis biological processes of lactylated proteins. (B) GO analysis of cellular components of lactylated proteins. (C) GO analysis of molecular functions of lactylated proteins. (D) KEGG analysis of lactylated proteins. (E) Scatter plot screening for candidate proteins.

To further understand how lactylated proteins affect FMDV replication, we submitted differentially modified proteins before and after infection to the STRING online database and used Cytoscape (version 3.10.2) to construct protein-protein interaction (PPI) networks. In the PPI network, betweenness centrality represents the measure of the number of shortest paths that pass through a node in the network. If a node lies on all pairs of nodes’ shortest paths, it will have the highest betweenness centrality value. This means that the nodes act as ‘bridges’ in the network, controlling communication and information exchange between different nodes. Degree centrality indicates the number of nodes that a given node is directly connected to. If a node has a high connectivity level with other nodes, its degree value will be correspondingly high. Through the determination of betweenness centrality and degree centrality, we identified seven key modified proteins, VDAC1, GAPDH, FAU, PALB2, EEF2, ATP5F1A, and RPS3. Among these proteins, VDAC1 has the highest values for both betweenness centrality and degree, indicating that it plays a crucial regulatory role in the overall network (Figure 6C).

### FMDV promotes VDAC1 lactylation and the release of mtDNA, activating the cGAS-STING signaling pathway

FMDV infection leads to the lactylation of many proteins in equine epithelial cells, but the role of this modification in virus replication requires further clarification. We first treated equine epithelial cells with 2.5 nM and 5.0 nM GSK2837808A and 10 mM lactate in the culture medium. The results of the immunoblotting experiment showed that when lactate dehydrogenase was inhibited, protein lactylation was also suppressed, and at this time, FMDV protein expression increased significantly (Figure 7A). When we added exogenous lactate to the porcine epithelial cells, we found that protein lactylation increased, and at this time, FMDV protein was significantly inhibited (Figure 7B), indicating that some proteins can inhibit virus replication after lactylation.

**Figure 7.**
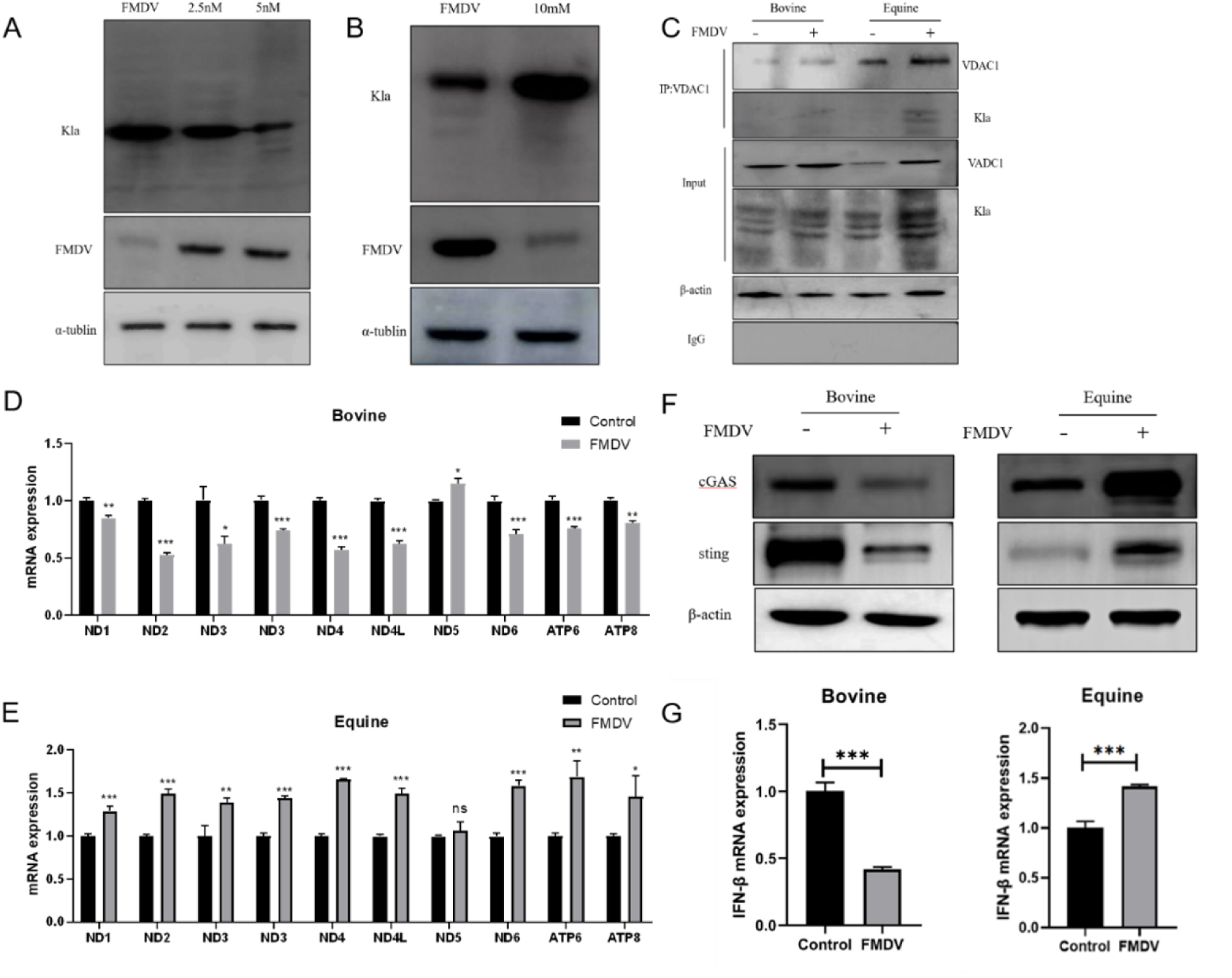
VDAC1 lactylation activates the cGAS-STING signaling pathway by inducing the release of mitochondrial DNA (mtDNA). (A) Effect of 2.5 nM and 5.0 nM GSK2837808A (lactate dehydrogenase inhibitor) on protein lactylation (Kla) and FMDV replication in porcine epithelial cells. (B) Effect of exogenous L-lactate addition on protein lactylation and virus replication in porcine epithelial cells. (C) Co-IP validation of VDAC1 lactylation. (D) FMDV infection of bovine epithelial cells suppresses the release of mtDNA into the cytoplasm. (E) FMDV infection of equine epithelial cells promotes the release of mtDNA into the cytoplasm. (F, G) Expression of IFN-β mRNA after FMDV infection. Experiments were repeated at least three times and values are means ± SD. Student’s *t*–test was used for determining the significance of differences: ns>0.05, \**p*< 0.05, \*\**p*< 0.01, \*\*\**p*< 0.001.

We previously found that VDAC1, a mitochondrial outer membrane channel protein, is located at the center of the protein network and is functionally important. Therefore, we determined whether VDAC1 was modified by lactylation and whether it affected FMDV replication. Co-IP experiments confirmed the interaction between VDAC1 and the pan-lactylation antibody, proving that VDAC1 was indeed lactylated after FMDV infection (Figure 7C).

VDAC1 controls the transport of materials between the mitochondria and the cytoplasm, including mtDNA, which depends on VDAC1 for entry and exit from the mitochondria. Therefore, we tested VDAC1 for FMDV-mediated lactylation and release of mtDNA. We detected an increase in the copy number of cytoplasmic mtDNA (cmtDNA) after FMDV infection. We selected mitochondrial DNA-encoded genes (ND1, ND2, ND3, ND4, ND4L, ND5, ND6, ATP6, ATP8) for qPCR quantification and normalized cmtDNA to total cellular mtDNA for analysis. The results showed that FMDV infection caused a significant decrease in cmtDNA copy number in bovine epithelial cells (*p*< 0.05) (Figure 7D), while in equine epithelial cells, cmtDNA copy numbers increased significantly except for ND5 (*p*< 0.05) (Figure 7E), indicating that FMDV stimulated the release of mtDNA from mitochondria into the cytoplasm in equine epithelial cells.

In published research on dengue virus (DENV), it was found that DENV caused mitochondrial damage, leading to the release of mtDNA into the cytoplasm and activation of the cGAS (cGMP-AMP synthase)-STING (stimulator of interferon genes) pathway for detection of cytosolic DNA. This research supports the idea that because FMDV is also an RNA virus, it could activate the cGAS-STING signaling pathway through the release of mtDNA. Subsequently, we examined the changes in the cGAS-STING signaling pathway in FMDV-infected epithelial cells. Our data showed that in FMDV-infected bovine epithelial cells, both cGAS and STING proteins exhibited significantly decreased expression levels, while in equine cells, the opposite was observed with both cGAS and STING being significantly upregulated by FMDV (Figure 7F). This suggests that after infection with FMDV, the modification of VDAC1 by lactylation in horses opens channels on the outer mitochondrial membrane, allowing mtDNA to pass into the cytoplasm and activate the cGAS-STING signaling pathway.

The activation of the cGAS-STING signaling pathway stimulates the production of type I interferon, which in turn activates the innate immune response. Here we showed that IFN-β release was significantly inhibited in cows infected with FMDV, while in horse epithelial cells, the expression of IFN-β increased significantly. Therefore, the resistance of horses to FMDV may be due to the inhibition of the virus by interferon produced after infection.

In a previous study, our research group used RNA-seq to compare the differences between bovine and equine nasopharyngeal tissues following FMDV infection, to explore the genetic basis of FMDV susceptibility at the transcriptional level. During bioinformatics analysis of the transcriptome data, we observed that upon FMDV infection the differentially expressed genes in the bovine nasopharynx were enriched in the apoptotic biological process (Figure 8A, B). Key genes promoting apoptosis, such as for caspase 3 (*CASP3*) and cytochrome C (*CYCS*), were upregulated in the transcriptomics data, and their expression was validated by RT-qPCR (Figure 8C).

**Figure 8.**
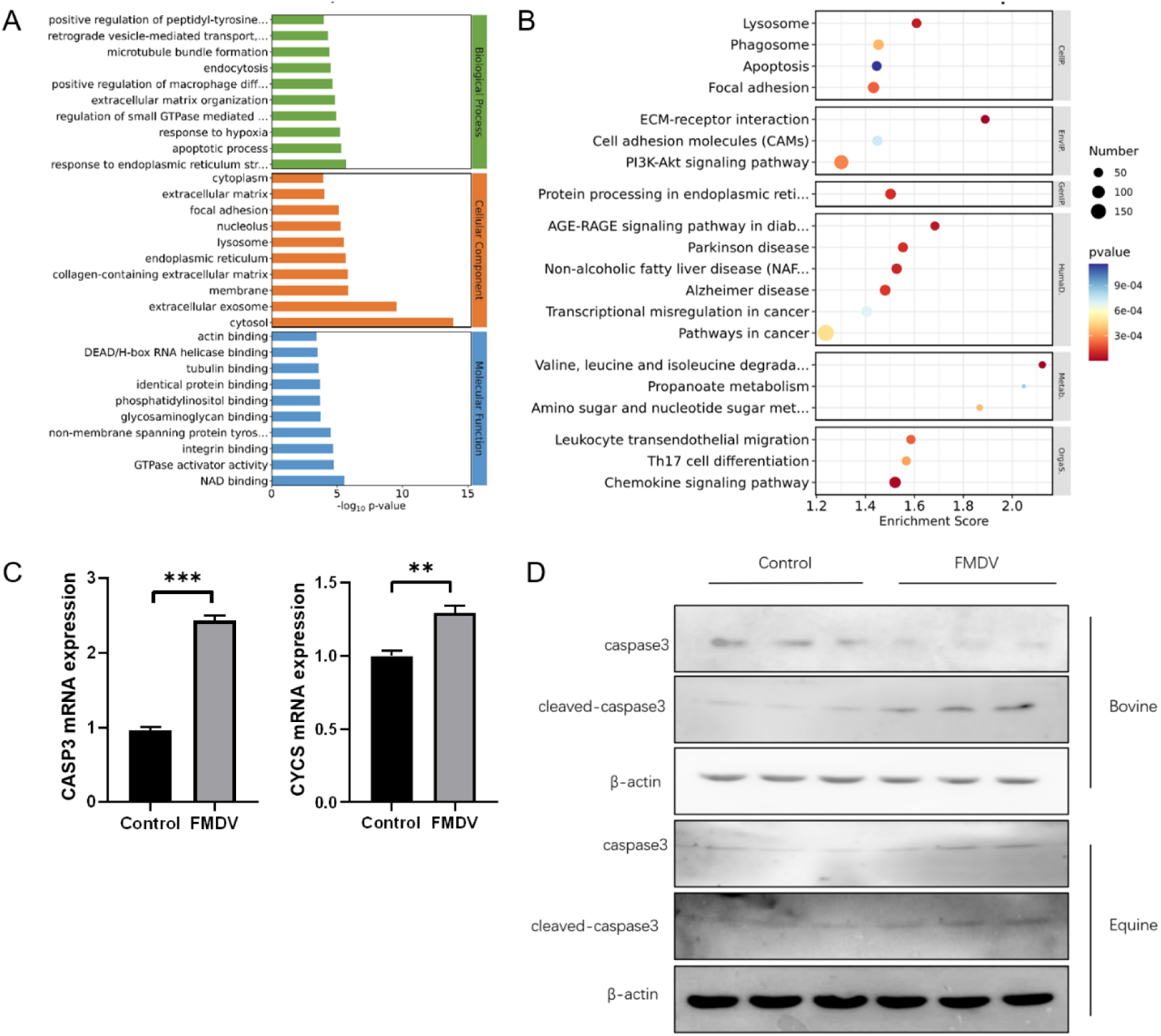
FMDV promotes apoptosis in bovine epithelial cells. (A) GO analysis of RNA-seq data. (B) KEGG analysis of RNA-seq data. (C) Expression of pro-apoptotic genes at the mRNA level. (D) Expression of apoptotic proteins in cows and horses before and after FMDV infection. All experiments were repeated at least three times and values are means ± SD. Student’s *t*–test for statistical significance was performed: \*\**p*< 0.01, \*\*\**p*< 0.001.

cGAS protein expression affects cGAS-mediated signaling pathways (Zheng, 2020; Lin and Xing et al., 2021), and various viruses have been found to regulate cGAS expression through caspase proteins. For example, ZIKV and vaccinia virus induce cGAS cleavage by caspase proteins associated with apoptosis (Zheng and Liu et al., 2018; Ning and Wang et al., 2019). Therefore, we further investigated the expression of caspase apoptotic proteins in FMDV-infected bovine epithelial cells. The results showed that intracellular caspase 3 protein decreased, while cleaved-caspase 3 increased significantly (Figure 8D), indicating that FMDV promoted apoptosis in bovine epithelial cells. The upregulation of apoptotic protein expression may be one of the reasons why the cGAS-STING signaling pathway is inhibited in bovine epithelial cells.

## Discussion

Our research group’s previous studies confirmed that horses and cattle share the same FMDV-targeting integrin receptor, which suggests that horses should be able to bind and internalize the virus. However, horses exhibit resistance to FMDV, leading us to speculate whether this was due to the activation of a distinct resistance mechanism in equine epithelial cells upon FMDV entry, compared with susceptible animals like cattle. Following FMDV infection, equine epithelial cells produced more lactic acid than bovine epithelial cells. The role of lactic acid in the resistance of horses to FMDV is a key aspect and we chose to investigate it further. Most viruses utilize the Warburg effect to provide energy and substrates for viral replication, leading to increased lactic acid production (Sanchez and Lagunoff, 2015). However, our data show that bovine and caprine epithelial cells did not exhibit the Warburg effect in response to FMDV infection but maintained normal oxidative phosphorylation activity. In contrast, the resistant animal, the horse, produced large amounts of lactic acid after FMDV infection. Some studies have found that lactic acid can enhance viral immune escape, but this does not seem to be the case in equine epithelial cells where lactic acid proved to have a strong inhibitory effect on FMDV replication. Our findings suggest that lactic acid plays a vital role in suppressing FMDV replication, making it a prime candidate for future research.

Lactylation is a novel emerging form of protein modification, which has provided new insights into many physiological processes. Most recent research has focused on the lactylation of histones; however, our group’s lactylation screening has shown that lactylation plays an essential role in the activity of proteins other than histones. The results of this study indicate that lactylation of the mitochondrial voltage-dependent anion channel 1 protein (VDAC1) has an inhibitory effect on viral replication, with significant differences observed in VDAC1 lactylation between bovine and equine epithelial cells.

VDAC1 is located on the outer mitochondrial membrane, which is a crucial organelle controlling various cellular functions, such as oxidative phosphorylation, thermogenesis, iron-sulfur cluster biogenesis, and biosynthesis of heme, lipids, and amino acids(Wallace, 2005)(Shoshan-Barmatz and De S et al., 2017; Shoshan-Barmatz and Krelin et al., 2017; Shoshan-Barmatz and Krelin et al., 2018). Mitochondrial dysfunction has been linked to numerous diseases and considered a causative factor in many neurodegenerative disorders, including Parkinson’s disease (PD), Huntington’s disease (HD), amyotrophic lateral sclerosis (ALS), and Alzheimer’s disease (AD) (Lezi and Swerdlow, 2012). The enrichment of Parkinson’s disease and Huntington’s disease signaling pathways in previous bioinformatics analyses suggests that foot-and-mouth disease virus (FMDV) may also induce changes in mitochondrial function.

The RIG-I-MDA5-mitochondrial antiviral signaling protein (MAVS) axis is the primary sensing pathway for RNA viruses, while the axis composed of cyclic GMP-AMP synthase (cGAS) and stimulator of interferon genes (STING) is the main sensing pathway for DNA viruses (Wu and Chen, 2014). cGAS is a cytoplasmic DNA sensor that primarily recognizes DNA released from the nucleus or mitochondria, or acquired from the extracellular microenvironment, including DNA from viral, retroviral, bacterial, and parasitic pathogens. Upon binding to a broad range of foreign and self dsDNA, cGAS catalyzes the synthesis of the second messenger cyclic GMP-AMP (cGAMP), which then binds to the adaptor protein STING located on the endoplasmic reticulum (ER). After dimerization, STING translocates from the ER to the Golgi apparatus, where it recruits Tank-binding kinase 1 (Tbk1), leading to the activation of the transcription factors NFκB and IRF3, which translocate to the nucleus to induce the production of type I IFN and activation of interferon-stimulated genes (ISGs).

cGAS, as a unique pathogen recognition receptor (PRR) for dsDNA, has received widespread attention in the study of DNA viruses. However, recent research has shown that RNA virus infections can activate the cGAS-STING signaling pathway by causing the release of mitochondrial DNA (mtDNA) into the cytoplasm(Li and Li et al., 2020), which promotes the production of type I interferon and induces an antiviral response. In our research, when horse cells were infected with FMDV, a large amount of mtDNA was released into the cytoplasm, and WB detection of cGAS and STING showed that the cGAS-STING signaling pathway in horses was promoted, which is the opposite of what was observed in cows. Therefore, horses might be unable to support rapid FMDV replication after infection because VDAC1 undergoes lactylation under metabolic regulation, which changes its oligomeric pore-forming structure in the OMM, thereby allowing the release of mtDNA into the cytoplasm. This activates the cGAS-STING signaling pathway, promoting the production of type I interferon and ultimately inhibiting viral replication. In contrast, both mtDNA release and the cGAS-STING signaling pathway in cows are inhibited during FMDV infection, the latter possibly due to viral activation of a series of caspase proteins that cleave cGAS, suppressing the cGAS-STING pathway, reducing interferon production, and facilitating viral replication by inhibiting innate immunity. In bovine cells infected with FMDV, the expression of caspase 3 decreased while cleaved-caspase 3 increased, indicating that FMDV promoted apoptosis in bovine epithelial cells. One reason why FMDV could replicate extensively in bovine epithelial cells may be due to apoptosis suppressing the cGAS-STING signaling pathway and reducing interferon production, creating a favorable environment for viral replication.

## Materials and Methods

### Cell culture, virus infection, and treatment with biochemical drugs

Using the tissue explant attachment method to culture nasopharyngeal epithelial cells, the nasopharyngeal tissue was rinsed with physiological saline to remove blood and mucus, and excess muscle was trimmed away from the tissue. The tissue was washed multiple times with PBS containing 1% antibiotics until the liquid was no longer cloudy, after which it was rinsed with 75% ethanol for 30 seconds, and washed three times with PBS containing 0.1% antibiotics. The edges of the cleaned tissue were removed and the central portions were placed into a vial with a small amount of DMEM/F12 complete medium (Gibco,Carlsbad,USA) containing 10% FBS (HyClone, Logan, UT, USA) and 0.1% antibiotics. The tissue was minced into small pieces and evenly spread onto a 25 cm² cell culture dish which was placed in a 37°C cell culture incubator for about 1 h to allow the tissue pieces to adhere tightly to the bottom. After the tissue pieces have adhered, slowly add 5 mL of complete medium and continue culturing, replacing the medium every three days. After a large number of cells have migrated out from the tissue pieces and formed a confluent layer, the pieces were rinsed away with PBS.

Under a microscope, mark areas with cells exhibiting obvious epithelial morphology, discard old medium and add a small amount of PBS to keep the bottom moist. With a sterile cotton swab, wipe off the cells outside the marked areas, then rinse with PBS to remove any debris. Add complete medium and continue culturing until the cells form a confluent monolayer, then subculture them. Pre-warm complete medium, PBS, and trypsin to 37°C. Discard the old medium, rinse the flask with pre-warmed PBS, then add 1 mL of trypsin and incubate at 37°C incubator for 4-6 minutes to loosen the cells. Add an equal volume of complete medium to stop the digestion, transfer the cell suspension to a 15 mL centrifuge tube and centrifuge at 1000 g for 5 minutes, then discard the supernatant. Resuspend the cell pellet in 1 mL of complete medium, count the cells, and seed them at a density of 1×10⁴ cells/mL with 5 mL of medium into a 25 cm² cell culture flask, and continue culturing in the incubator. Resuspend the remaining cells in freezing medium, transfer to a cryovial, place the cryovial in a programmed freezing container, and store it at −80°C for 24 hours. The next day, transfer the cryovial to liquid nitrogen for long-term storage.

The cells were infected with the FMDV/OHC/02 strain (Bioway AnTai Biotechnology Co., Ltd). Mix the virus suspension with basal medium at a ratio of 1:4, then add to the cell culture flask for virus infection. GSK2837808A and L-lactate (Solarbio, Beijing, China) were dissolved in dimethyl sulfoxide (DMSO), then diluted in DMEM/F12 medium containing 10% FBS and applied to the cells at final concentrations of 2.5 nM and 5 nM for GSK2837808A, and 10 mM for L-lactate.

### Real-time quantitative PCR (RT-qPCR) for relative transcript levels

Total RNA was extracted using a TRIzol reagent kit according to instructions. The concentration and purity of the RNA were measured using a NanoDrop 2000c nucleic acid analyzer, then 500 ng of each RNA sample was taken for reverse transcription. A 10 μL reaction was prepared using the PrimeScript™ reagent kit, with the reverse transcription conditions set as follows: 37°C for 15 minutes, 85°C for 5 seconds, and hold at 4°C.

The SYBR Premix Ex Taq™ kit was used to prepare a 20 μL reaction for RT-qPCR. according to the kit instructions, and amplification was performed on an ABI7500 real-time PCR instrument. The amplification conditions were: 95°C for 30 seconds, then 40 cycles of 95°C for 5 seconds, 60°C for 34 seconds; and lastly, one cycle of 95°C for 15 seconds, 60°C for 1 minute, and 95°C for 15 seconds. Each reaction was repeated three times, and dissociation curves were analyzed after each run to verify target specificity.

The relative mRNA expression levels were calculated using the 2^-ΔΔCT^ method, with *ACTB* as the internal control.

### Antibody Microarray Preparation

After multiple injections of mice with MYA98, we isolated B lymphocytes, which were then fused with cultured myeloma cells. Through limiting dilution, positive hybridoma clones were identified, and the monoclonal cells were expanded by cultivation. The IgG antibodies in cell supernatants were then purified by binding to protein G agarose beads. The collected antibodies were spotted onto the chip carrier (FAST chip, CapitalBio, City, Country) using an inkjet spotting instrument (Arrayjet Ltd., Roslin, UK), with each antibody spotted at 10 nL. After chip fabrication, quality control was performed.

### Western blotting

Total proteins were extracted from cell samples using RIPA lysis buffer containing PMSF and a phosphatase inhibitor cocktail (Solarbio, Beijing, China). Protein concentrations were determined using the Pierce BCA protein assay kit (Thermo Fisher Scientific, Germany). Equal amounts of proteins were separated by SDS-PAGE on an 8% polyacrylamide gel and then transferred to a PVDF membrane (Millipore, USA). The membrane was blocked with 5% skim milk in TBST buffer at room temperature (RT) for 3 h. After blocking, the membrane was incubated with appropriately diluted primary antibodies at 4°C overnight, washed in TBST, and incubated with horseradish peroxidase (HRP)-conjugated goat anti-rabbit IgG (1:5000 dilution) at RT for 1 h. Immunoreactive proteins were visualized using an ECL staining kit (Beyotime, Shanghai, China), images were captured using an imaging system (FluorChem Q, USA), and band densities were calculated. The primary antibodies used in this study included anti-PFKP (7C10), anti-PKM (Cell Signaling Technology, Danvers, MA, USA), anti-HK1, anti-cGAS, anti-STING, anti-VDAC1, anti-Pan-Kla (Jingjie, Hangzhou, China), anti-caspase-3, anti-cleaved caspase-3, anti-LDHA (Proteintech, USA), and anti-β-actin (Abcam, USA).

### Extracellular acidification rate (ECAR) assay

The P61 probe (stock solution) was diluted 10-fold using reagent B assay buffer (900 μL of assay buffer to 100 μL of P61 probe). The culture medium was removed from the cells to be tested, the cells were washed twice with assay buffer, and 90 μL of fresh assay buffer was added to each well together with 10 μL of BBcellProbe® P61 acidification fluorescence probe. A microplate reader was used to detect changes in extracellular acidification rate by measuring the change in fluorescence over time. The excitation wavelength was 488 nm and the emission wavelength was 580-590 nm. The plate was read every 2-3 min, with a total measurement time not exceeding 120 min. The acidification rate curve was plotted for each sample and the slope was calculated.

### Oxygen consumption rate (OCR) assay

The culture medium was removed from the cells to be tested, they were washed with PBS, 100 μL of fresh medium was added to each well along with 4 μL of BBoxiProbe® R01 oxygen fluorescence probe and 100 μL of oxygen sealing solution. The plate was read every 2-3 min using a fluorescence plate reader with excitation wavelength of 455-468 nm and emission wavelength of 603 nm to measure changes in extracellular O2 consumption.. The oxygen consumption rate (OCR) curve was plotted and the slope of each sample curve was calculated.

## Acknowledgments

We are grateful for the support of the virus laboratory facility provided by the Inner Mongolia Bit Bio-Tech Co., Ltd.

